# Cytolethal distending toxin promotes epithelial-to-mesenchymal transition by modulating AKT-Dependent β-catenin Ser552 phosphorylation

**DOI:** 10.64898/2026.06.01.729203

**Authors:** Ruxue Jia, Lamia Azzi-Martin, Mariana Saraiva, Elodie Sifré, Christine Varon, Pierre Dubus, Armelle Ménard

**Author notes:** Corresponding author: Dr. Armelle Ménard, Université de Bordeaux, Campus de Carreire, Bâtiment BBS, BoRdeaux Institute of onCology - BRIC INSERM UMR1312, F-33000, Bordeaux, France –.

## Abstract

Bacterial genotoxins, Cytolethal Distending Toxin (CDT) and colibactin, cause DNA damage in intoxicated epithelial cells of the host. DNA damage influences β-catenin signaling, altering its stability and nuclear translocation, potentially contributing to cancer development. Using non-transformed hepatocytes and cancer-derived intestinal and hepatic epithelial cell lines, we showed that CDT/CdtB induces the phosphorylation of β-catenin at serine 552, along with the loss of β-catenin from adherens junctions. This leads to the subsequent cytoplasmic accumulation and nuclear translocation of β-catenin, ultimately driving TCF/LEF transcription, the crucial downstream event of Wnt/β-catenin signaling, as well as the transcription of some β-catenin target genes. Colibactin induces similar effects. MK-2206, a direct AKT inhibitor, and metformin, an AMP-activated protein kinase activator that indirectly inhibits AKT, both protected cells against various effects induced by CdtB exposure. These effects include β-catenin phosphorylation at Ser552, the disassembly of cell-cell junctions and the subsequent nuclear accumulation of phosphorylated β-catenin, leading to reduced TCF/LEF-mediated transcription. Additionally, MK-2206 and metformin protected from CdtB-induced epithelial–mesenchymal transition (EMT), including increased nuclear accumulation of SNAIL, enhanced matrix degradation and motility. Overall, these data show that infection with genotoxin-producing bacteria controls some EMT features through β-catenin and AKT-dependent signaling.

## Introduction

Certain bacteria that produce colibactin and cytolethal distending toxin (CDT) are associated with several cancers, particularly colorectal cancer (CRC). By damaging host DNA, these toxins induce genomic instability and heritable mutations that potentially drive carcinogenesis [1,2]. Beyond DNA damage, CDT and colibactin facilitate malignancy by promoting polyploidy [3], inflammatory responses [4,5], anchorage-independent growth [1], while simultaneously targeting pathways essential for cell transformation [1,6–9]. Evidence from both murine and human studies supports a significant role for these toxins in cancer development [10–14]. We have shown that infection with genotoxin-producing bacteria elicits epithelial to mesenchymal transition (EMT) [15], a crucial process in cancer initiation and/or progression by which cells lose their epithelial characteristics to gain mesenchymal properties conducive to cell migration and invasion. Both colibactin and CDT induced the disassembly and removal of junctional proteins from the plasma membrane at tight, adherens and desmosomal junctions [15].

The interplay between E-cadherin and β-catenin is crucial in maintaining epithelial integrity. Under normal conditions, E-cadherin binds to β-catenin at the cell plasma membrane, strengthening cell adhesion (keeping the adherens junctions intact), and preventing uncontrolled cell growth by sequestering β-catenin away from the nucleus [16]. However, when E-cadherin undergoes shedding, β-catenin loses its membrane-bound anchor. This free cytosolic β-catenin can translocate to the nucleus, acting as a transcriptional co-activator, ultimately promoting genes involved in cell proliferation, differentiation, and survival [16]. The Wnt/β-catenin signaling pathway is important in development, wound healing, and can also contribute to cancer progression if not tightly controlled.

As the primary downstream effector of Wnt signaling, β-catenin serves as both a structural component of cell junctions and a transcriptional co-activator within the nucleus. Its subcellular location and transcriptional activity are tightly regulated by multiple phosphorylation events. Phosphorylation in the N-terminal region (at Ser33, Ser37, and Thr41) by GSK3β targets β-catenin for degradation, whereas other phosphorylations increase its stability and enhance its activity [17], notably at Ser191/605 by JNK2 (c-Jun N-terminal kinase 2/ Mitogen-Activated Protein Kinase 9) [18], Ser552 by AKT [19], Ser675 by PKA/PRKACA (Protein Kinase A/Protein Kinase cAMP-Activated Catalytic Subunit Alpha) [20]. Phosphorylation at Ser552 by AKT kinase (also known as PKB for protein kinase B) is a widely recognized event that promotes β-catenin’s dissociation and release from cell-cell contacts [19]. This leads to its accumulation in both the cytosol and nucleus. In the nucleus, β-catenin primarily interacts, through its C-terminal transactivation domain, with T-cell factor/lymphoid enhancer-binding factor (TCF/LEF) transcription factors, acting as a transcriptional co-activator, to enhance transcriptional programs [19].

Since CDT and colibactin trigger a loss of cell-cell junctions [15], this is expected to result in increased cytoplasmic levels of free β-catenin, potentially leading to its nuclear translocation and the subsequent activation of β-catenin target genes. Accordingly, the present study aimed to evaluate the role of bacterial genotoxins in modulating β-catenin expression, subcellular localization and activity. These analyses were performed on non-transformed hepatocytes, and cancer-derived intestinal and hepatic epithelial cell lines following infection with *H. hepaticus* or its corresponding CDT-knockout mutant strain. The effects were attributed to the active CdtB subunit of CDT following its ectopic production. To ensure that the observed effects could be specifically attributed to CdtB, a corresponding mutated CdtB lacking catalytic activity was used as a control. Finally, this study was extended to colibactin, a genotoxic secondary metabolite produced by *Escherichia coli* [21].

## Results

### *Helicobacter hepaticus* infection alters β-catenin localization

The effect of *H. hepaticus* infection on β-catenin subcellular localization was analyzed in the liver of mice either infected or uninfected, for 14 months [22]. Immunohistochemical analyses of the liver of non-infected mice showed that β-catenin was primarily localized at cell-cell junctions rather than in the nuclei (Fig. 1A). In mice infected with *H. hepaticus*, the infection primarily leads to a moderate increase in the overall level of nuclear β-catenin within cells with preserved cell-cell junctions (Fig. 1A, *Hh*_L_, 1.5-fold increase relative to non-infected mice). However, in certain areas (Fig. 1A, *Hh*_H_), β-catenin staining disappeared from cell-cell junctions, increased in the cytoplasm and accumulated in the nuclei of hepatocytes, resulting in a significant increase in nuclear β-catenin levels (2.9-fold increase compared to non-infected mice). This suggests that *H. hepaticus* infection alters the subcellular localization of β-catenin in the liver of mice.

**Figure 1.**
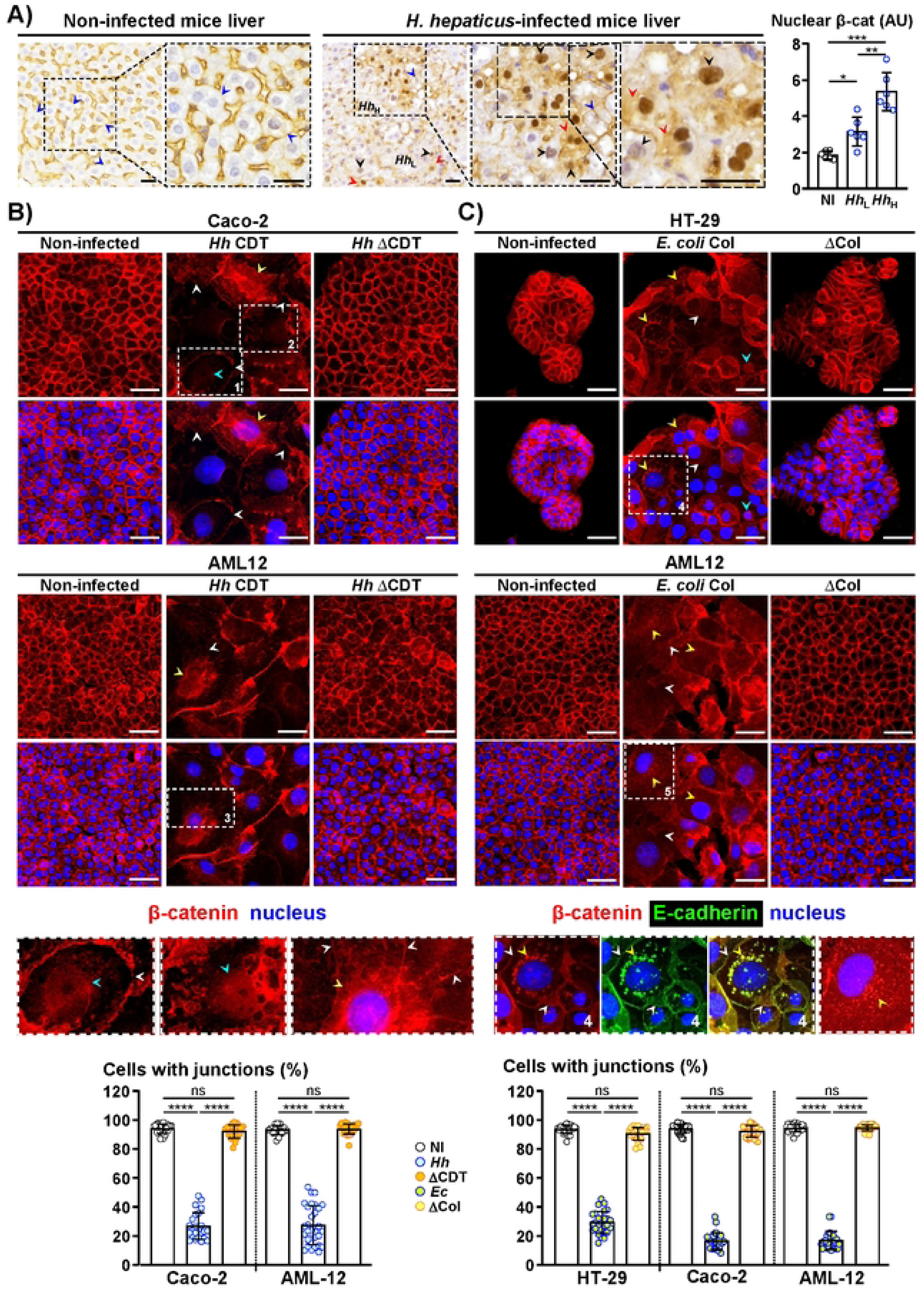
Infection with genotoxin-producing bacteria alters β-catenin localization. **A)** Images of non-transgenic mouse livers, either uninfected or infected with *H. hepaticus* for 14 months, immunostained for β-catenin and counterstained with hematoxylin. Magnifications of selected areas are shown in boxes. Scale bar, 100 μm. Blue arrowheads indicate junctional β-catenin. Black arrowheads indicate variable intensities of nuclear β-catenin. Red arrowheads indicate cytoplasmic accumulation of β-catenin. *Hh*_L_ and *Hh*_H_ represent areas within the livers of mice infected with *H. hepaticus* mice that exhibit low and high levels of nuclear β-catenin, respectively. Protein nuclear intensity was quantified. Data are presented as the mean ± SD (n=6 mice). *p<0.05, **p<0.01, and ***p<0.001 (n=6, ANOVA and Tamhane’s T2 post-hoc test). **B)** and **C)** Cells were infected with *H. hepaticus* producing CDT, or *E. coli* producing colibactin or their corresponding knock-out mutant lacking genotoxic activity, ΔCDT or ΔCol, and uninfected control. At 96 hours post-infection, cells were probed with primary antibodies generated against cell-cell junctions followed by fluorescently labeled secondary antibodies (β-catenin: red; E-cadherin: green) and counterstained with DAPI to highlight the nuclei (blue). A more intense color and higher magnification of selected areas are shown in boxes. Scale bar, 50 μm. White arrowheads indicate the loss of junctional β-catenin and residual membrane spans containing β-catenin in intoxicated cells. Yellow arrowheads indicate the cytoplasmic accumulation and perinuclear relocalization of β-catenin. Relocalization of E-cadherin from the plasma membrane to the perinuclear region indicates a loss of functional cell-cell adherens junctions. This shift is a hallmark of EMT, reflecting E-cadherin trafficking and ubiquitination, followed by degradation *via* the perinuclear recycling endosomes—a process essential for the transition toward a migratory phenotype. Blue arrowhead indicate nuclear accumulation of β-catenin. Cells with junctions were manually counted across 3 independent experiments; each utilized triplicate coverslips divided into three sections. Data represent the mean ± SD (n=27). ****p<0.0001 (Kruskal-Wallis and Dunn’s post-hoc test). The vertical lines separate the analyses performed on different cell lines. **B)** Images of cells either uninfected or infected with *H. hepaticus* (strain 3B1/Hh-1) or its corresponding ΔCDT mutant. Since HT-29 cells are resistant to the effects of CDT during infection experiments, they were not used. **C)** Images of cells either uninfected or infected with *E. coli*, either producing colibactin or not. *Abbreviations:* AU, arbitrary units; β-cat, β-catenin; CDT, cytolethal distending toxin; Col, colibactin; DAPI, 4ʹ,6-diamidino-2-phenylindole; *Ec, E. coli* producing colibactin; *Hh*, *H. hepaticus*; *Hh*_L,_ and *Hh*_H,_ areas in *H. hepaticus* infected mice with low and high levels of nuclear β-catenin; NI, non-infected; ns, not significant; ΔCDT, CDT-knockout mutant strain; ΔCol, *E. coli* not producing colibactin.

CDT, being the main virulence factor of *H. hepaticus*, its effect on adherens junctions was evaluated by immunofluorescence on epithelial intestinal and hepatic cell lines (Fig. 1B), as this bacterium colonizes both sites [23]. Murine non-transformed AML12 (hepatic) cells and human cancer-derived cell lines—Caco-2 (intestinal) and HT-29 (intestinal)—were infected with CDT-producing bacteria, except for HT-29 cells, which are resistant to the effects of CDT during infection experiments with *Helicobacter* species [23]. In non-infected cells, β-catenin was mainly detected at the plasma membrane and showed well-delineated adherens junctions. Upon *H. hepaticus* infection, β-catenin staining disappeared from cell-cell junctions (white arrowheads, enlargement in boxes 1 and 3), with sometimes residual membrane spans, and accumulated in the cytoplasm and around the nucleus (yellow arrowheads, enlargement in boxes 3 and 5). A nuclear accumulation of β-catenin was also observed in some intoxicated cells (blue arrowheads, enlargement in boxes 1 and 2). The disassembly of cell-cell junctions led to approximately 27% of Caco-2 and AML12 infected cells retaining cell-cell junctions (Fig. 1B). These effects were not observed in cells infected with the CDT-knockout mutant strain lacking CDT activity, suggesting that CDT is likely the causative factor.

### CDT/CdtB promote the loss of adherens epithelial cell junctions

The involvement of CDT was confirmed upon ectopic production of the CdtB subunit of *H. hepaticus* and the corresponding inactive CdtB_mut_ [23]. Stable transgenic cell lines (Fig. S1), including HT-29 cells—which are susceptible to CdtB when it is expressed intracellularly [23]—were used to enable the conditional ectopic production of *H. hepaticus* CdtB and its catalytically inactive mutant, CdtB_mut_ [24]. In Hep3B cells, cell-cell junctions exhibit weak cohesion and appear diffuse, which hinders efficient quantification [15]. However, upon CdtB exposure, residual membrane spans were observed along with a perinuclear accumulation of β-catenin puncta in some CdtB-intoxicated Hep3B cells (Fig. S1A, boxes 5 and 6, white and yellow arrowheads, respectively). HT-29 cells, Caco-2 and AML12 cells showed cell-cell junction disassembly, with only 24, 16 and 34% of cells retaining junctions following CdtB exposure (Fig. S1A). Additionally, β-catenin was found to accumulate in the nuclei of certain cells exposed to CdtB (Fig. S1A, boxes 1-3). β-catenin is a central component of the E-cadherin complex, and its redistribution is inherently tied to the integrity of adherens junctions. Consistent with this, the loss of β-catenin at cell-cell junctions corresponded to the loss of E-cadherin (Fig. S1B).

### Colibactin promotes the loss of adherens epithelial cell junctions

Colibactin-producing bacteria, such as *Klebsiella pneumonia* and *E. coli*, can colonize the gastrointestinal tract and disseminate to the liver [25,26]. Intestinal and liver cells were infected with *E. coli*, either producing colibactin or not (Fig. 1C). Infection with colibactin-producing *E. coli* also induced the disassembly and removal of β-catenin from adherens junctions, with approximately 29.2%, 16.5% and 16.9% of HT-29, Caco-2 and AML12 infected cells retaining cell-cell junctions, respectively. This was accompanied by a cytoplasmic and perinuclear accumulation of β-catenin puncta in some intoxicated cells (Fig. 1C, boxes 4 and 5). These effects were not observed in uninfected cells nor in cells infected with *E. coli* hosting the bacterial artificial chromosome vector (*pks-*) that does not express colibactin, indicating that the observed effects were induced by colibactin. The destabilization of the E-cadherin/β-catenin complex was also observed in colibactin-intoxicated cells (Box 4, Fig. 1C). The subsequent delocalization of E-cadherin from the plasma membrane to the perinuclear compartment marks a loss of functional adherens junctions. This shift is a hallmark of EMT, driven by E-cadherin trafficking and ubiquitination, followed by degradation or recycling *via* perinuclear endosomes—a process essential for the transition toward a migratory phenotype [27].

Taken together, these data show that CDT/CdtB and colibactin trigger a significant loss of cell-cell junction integrity with the disassembly and removal of existing adherens cell junctions.

### CdtB induces nuclear accumulation of phosphorylated β-catenin at Ser552

Phosphorylation of β-catenin at residue 552 promotes its dissociation from cell-cell junctions and accumulation in the cytosol and nucleus [19]. Western blot and immunocytofluorescence analyses were performed to localize and quantify β-catenin and its phosphorylated form at Ser552 following exposure to ectopic CdtB and CdtB_mut_. Compared to CdtB_mut_, the level of β-catenin remained unchanged upon CdtB exposure, while the phosphorylated form of β-catenin at Ser552 accumulated in CdtB-intoxicated cells (Fig. 2A, white bars). The ratio of [phospho-β-catenin-Ser552/total β-catenin] increased following CdtB exposure for all cell lines (Fig. 2A, grey bars), demonstrating an increase in phospho-β-catenin-Ser552, which should subsequently translocate to the nucleus to enhance transcriptional programs. In addition, CdtB exposure did not led to the phosphorylation of β-catenin in its N-terminal region at Ser33-37, except for HT-29 cells (Fig. S3), suggesting concomitant β-catenin degradation (cytoplasm) and activation (nucleus) in HT-29 cells.

**Figure 2.**
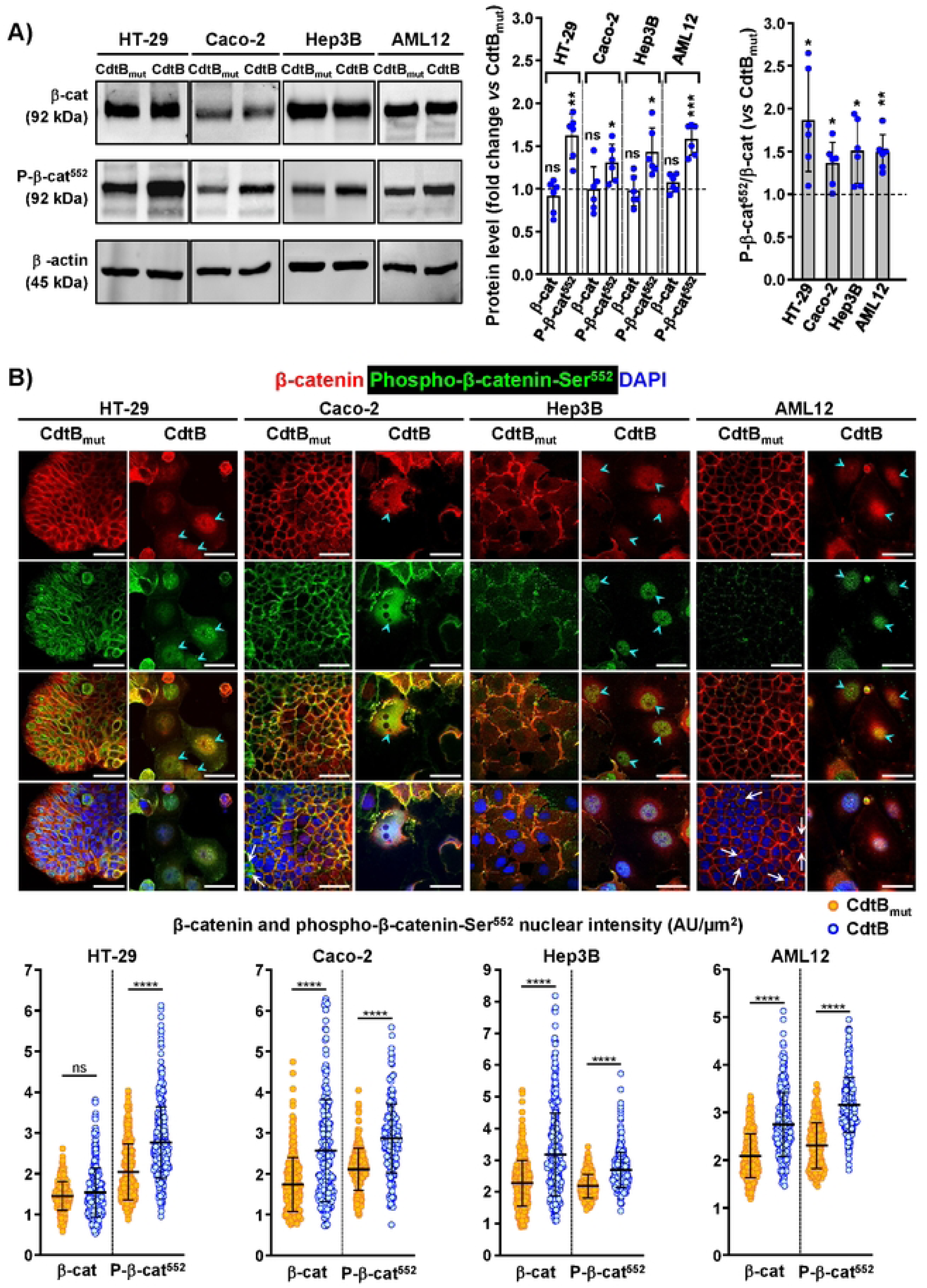
CdtB enhance nuclear accumulation of β-catenin phosphorylated at Ser552. Transgenic CdtB and CdtB_mut_ HT 29, Caco 2, Hep3B and AML12 cells were cultured in the presence of doxycycline, to induce the expression of CdtB or its corresponding inactive mutant, CdtBmut, respectively. The analyses were conducted 96 hours after induction. **(A)** Western blot analysis of the protein level of β-catenin and its phosphorylated form at Ser552. Quantification was conducted relative to total protein levels, using stain-free blots for normalization. β-actin is also presented. Data are presented as the mean ± SD (n=6). *p<0.05, **p<0.01, and ***p<0.001 relative to CdtB_mut_ (t-test). The horizontal dashed lines shows the basal rate of protein accumulation in cells exposed to CdtB_mut_. The vertical lines separate the analyses performed on different cell lines. Grey bars represent the [P-βcat-Ser552/total β-cat] ratio in CdtB-treated cells, normalized to the CdtB_mut_ control (set to 1). **(B)** Confocal images (z-slice) showing β-catenin (red) and its phosphorylated form at Ser552 (green). Cells were probed with primary antibodies directed against β-catenin (red) and its phosphorylated form at Ser552 (green) followed by fluorescently labeled-secondary antibodies and DAPI to counterstain the nuclei (blue). Scale bar, 50 µm. Blue arrowhead indicate nuclear accumulation of β-catenin. White long arrows (in merge) indicate bright, condensed chromatin in cells undergoing mitosis. Protein nuclear intensity was quantified. Data are presented as the mean ± SD (n=3, >300 nuclei). ****p<0.0001 relative to CdtB_mut_ (Mann-Whitney U test). Confocal microscopy projections of CdtB-intoxicated cells are presented in Supplemental Fig. S2A, illustrating Ser552-phosphorylated β-catenin nuclear accumulation. Additional wide-field images providing a clearer visualization of cell-cell junctions are provided in Supplemental Fig. S2B. *Abbreviations*: AU, arbitrary units; β-cat, β-catenin; CdtB, cytolethal distending toxin active subunit B of *H. hepaticus* strain 3B1; CdtB_mut_, mutated CdtB lacking catalytic activity; ns, non-significant; P-β-cat^552^, β-catenin phosphorylated at Ser552; Pser552.

Immunocytofluorescence analyses were performed to localize and quantify β-catenin and its phosphorylated form at Ser552. In contrast to CdtB_mut_ expressing cells, which retained organized adherens junctions (Fig. 2B and S2A, except for Hep3B), cells producing active CdtB exhibited junctional disassembly alongside elevated nuclear phospho-β-catenin-Ser552 (Fig. 2B and S2A, Hep3B included). This finding was further confirmed by confocal analysis (Fig. 2B and S2B).

### Exposure to CdtB, Colibactin, and DNA-damaging agents activates β-catenin-mediated transcription

Active nuclear β-catenin primarily interacts with TCF/LEF transcription factors, thereby activating specific transcriptional programs. The genes *AXIN2*, *MYC*, *CCND1*, *LGR5*, and *BMP4* are well-established direct target genes of TCF/LEF [28]. Their expression was assessed by RT-qPCR (Fig. 3A). CdtB induced increased expression of *AXIN2*, *CCND1* and *LGR5* genes in HT-29, as well as that of *AXIN2*, *MYC* and *LGR5* in Hep3B. Caco-2 cells showed a slight CdtB-downregulation of *MYC* and *LGR5* transcripts, concurrent with an upregulation of *AXIN2, BMP4* and *CCND1* transcripts. In non-transformed AML12 hepatocytes, *AXIN2* was not amplified; CdtB downregulated *LGR5* and *BMP4* genes and upregulated *CCND1* gene. Since β-catenin target gene expression differs among various cell lines, and as TCF/LEF is considered the primary transcriptional partner of β-catenin, we conducted a TCF/LEF luciferase reporter assay to assess β-catenin-dependent transcriptional activity [29]. High TCF/LEF transcriptional activity was found in naïve HT-29 and Caco-2 cells, while lower levels were found in naïve Hep3B and AML12 hepatocytes (Fig. 3B). These data are consistent with previous studies showing that cell lines derived from a colon adenocarcinoma have constitutive activation of the Wnt/β-catenin pathway [30–32], and that Hep3B and AML12 hepatocytes normally exhibit moderate to low levels of Wnt/β-catenin signaling activity [33]. Caco-2 cells exhibited 10-fold higher basal activity than other cell lines (Fig. 3B). Caco-2 cells harbor a mutation in *CTNNB1* gene, resulting in constitutive activation of the β-catenin signaling pathway [30,32]. Following exposure to CdtB, a moderate decrease in β-catenin transcription was detected in Caco-2 cells (Fig. 3C). On the other hand, a significant increase in β-catenin activity was observed in HT-29, Hep3B, and AML12 cells exposed to CdtB.

**Figure 3.**
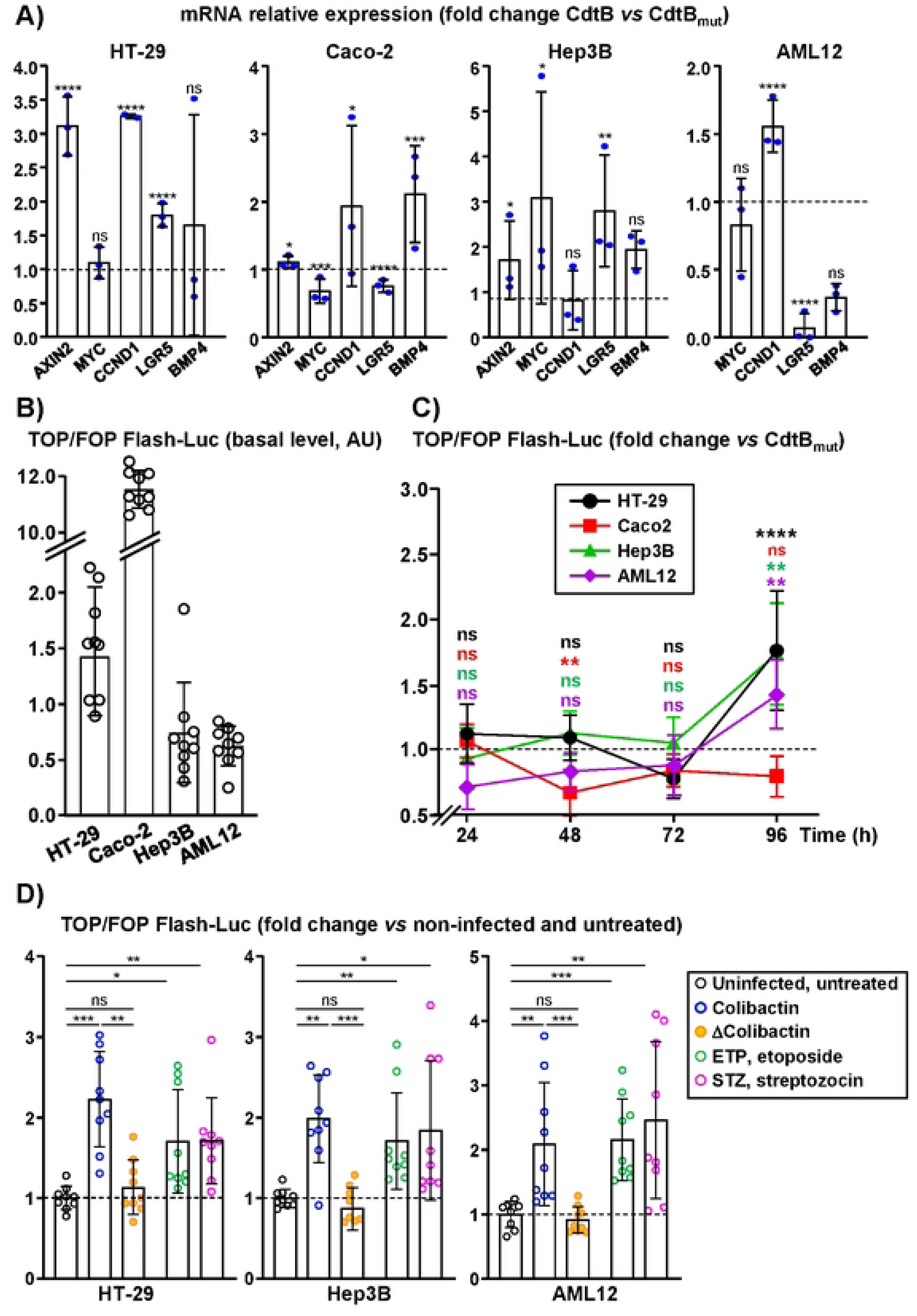
Exposure to CdtB, colibactin, and DNA damaging agents increases β-catenin transcriptional activity. **A)** Transgenic HT-29, Caco-2, Hep3B and AML12 cells were cultured in the presence of doxycycline, to induce the expression of CdtB or CdtB_mut_. At 96 hours post-induction, the relative expression of genes in cells was measured by RT-qPCR and normalized according to the expression of the reference gene, hypoxanthine phosphoribosyltransferase 1. Ratios were calculated using the 2^−ΔΔCt^ method. The relative gene expression in response to CdtB is reported as a fold change compared to cells exposed to CdtB_mut_. Results are presented as means ± SD of 3 independent experiments, each performed in triplicate. *p<0.05, **p<0.01, ***p<0.001 and ****p<0.0001 relative to CdtB_mut_ (n=3, t-test). The dashed line shows the basal rate of gene expression in cells exposed to CdtB_mut_. **B)** TOPflash/FOPflash-luciferase reporter activity of naïve HT-29, Caco-2, Hep3B and AML12 cells (basal level of β-catenin activity in cells non-exposed to bacterial genotoxin). Results are presented as means ± SD of 3 independent experiments, each performed in triplicate (n=9). **C)** Kinetics of TOPflash/FOPflash-luciferase reporter activity of transgenic cells cultured in the presence of doxycycline, to induce the expression of CdtB and CdtB_mut_. The dashed line shows the basal rate of gene expression in cells exposed to CdtB_mut_. Results are presented as means ± SD of 3 independent experiments, each performed in triplicate. ****p<0.0001 relative to CdtB_mut_ (ANOVA and Tamhane’s T2 post-hoc test). **D)** TOPflash/FOPflash-luciferase reporter activity assessed in naïve HT-29, Hep3B, and AML12 cells following infection with *E. coli* (either producing colibactin or not) or exposure to the DNA-damaging agents streptozocin (5 mM) and etoposide (2 μM). Activity was analyzed 96 hours post-treatment. The horizontal dashed line represents the basal gene expression level in uninfected and untreated cells. *p<0.05, **p<0.01, and ***p<0.001 (n=9, ANOVA and Tamhane’s T2 post-hoc test). *Abbreviations:* CdtB, cytolethal distending toxin active subunit B; CdtB_mut_, mutated CdtB lacking catalytic activity; Col, *E. coli* producing colibactin; ns, not significant; ΔCol, *E. coli* not producing colibactin.

Infection with colibactin-producing *E. coli* increased TCF/LEF transcriptional activity leading to an approximate 2-fold increase for HT-29, Hep3B and AML12 cells (Fig. 3D). These effects were not observed in uninfected cells and in cells infected with *E. coli* not expressing colibactin, demonstrating that the observed effects were induced by colibactin.

Both CDT and colibactin induce DNA damage, suggesting that their effects on β-catenin signaling are primarily due to the DNA damage they cause. To further explore this relationship, we evaluated DNA damaging agents with different mechanisms of action. We found that both etoposide, a topoisomerase II enzyme inhibitor, and streptozocin, a glucosamine-nitrosourea alkylating compound, enhanced TCF/LEF transcriptional activity (Fig. 3D). Thus the genotoxic stress induced by bacterial genotoxins activates β-catenin/TCF/LEF-mediated transcription.

### MK-2206 and metformin attenuate phosphorylation of β-catenin at Ser552 and CdtB-induced TCF/LEF-mediated transcription

AKT is the primary kinase mediating phosphorylation of β-catenin at Ser552. The effects of a highly selective allosteric inhibitor of panAKT (AKT1, AKT2, AKT3), MK-2206, and a non-selective inhibitor of AKT, metformin, were evaluated on HT-29, Hep3B and AML12 cells. All three naïve cells were sensitive to 48 h treatment with MK-2206, exhibiting dose-dependent inhibition of cell growth (Fig. S4). The concentration of MK-2206 was set at 2 µM. Metformin exhibited minimal toxicity in HT-29 and Hep3B cells, so its concentration was set at 5 mM. However, for AML12 cells, a lower concentration of 1 mM was selected for subsequent analyses. At the chosen concentrations, treatment with MK-2206 and metformin did not affect the increase in γH_2_AX accumulation, a marker of DNA damage (Fig. S5), nor the nuclear remodeling induced by CdtB (Fig. S6).

Western blot analysis (Fig. 4A) of the total cellular levels of β-catenin, AKT, and AMPK revealed that the overall levels of these proteins remained unchanged following exposure to CdtB_mut_ or CdtB in both cancer-derived HT-29 and Hep3B cell lines, as well as in non-cancerous AML12 cells. However, CdtB exposure led to significant changes in the phosphorylation of these proteins. Specifically, it increased β-catenin phosphorylation at Ser552 in HT-29, Hep3B, and AML12 cells (by 1.87-, 1.65-, and 1.88-fold, respectively). It also enhanced AKT phosphorylation at Ser473 in HT-29, Hep3B, and AML12 cells (by 1.75-, 1.79-, and 1.42-fold, respectively). Additionally, CdtB exposure resulted in AMPK phosphorylation at Thr172 in HT-29 and Hep3B cells, with increases of 2.82- and 1.71-fold, respectively, while only a tendency toward increased phosphorylated AMPK was observed in AML12 cells.

**Figure 4.**
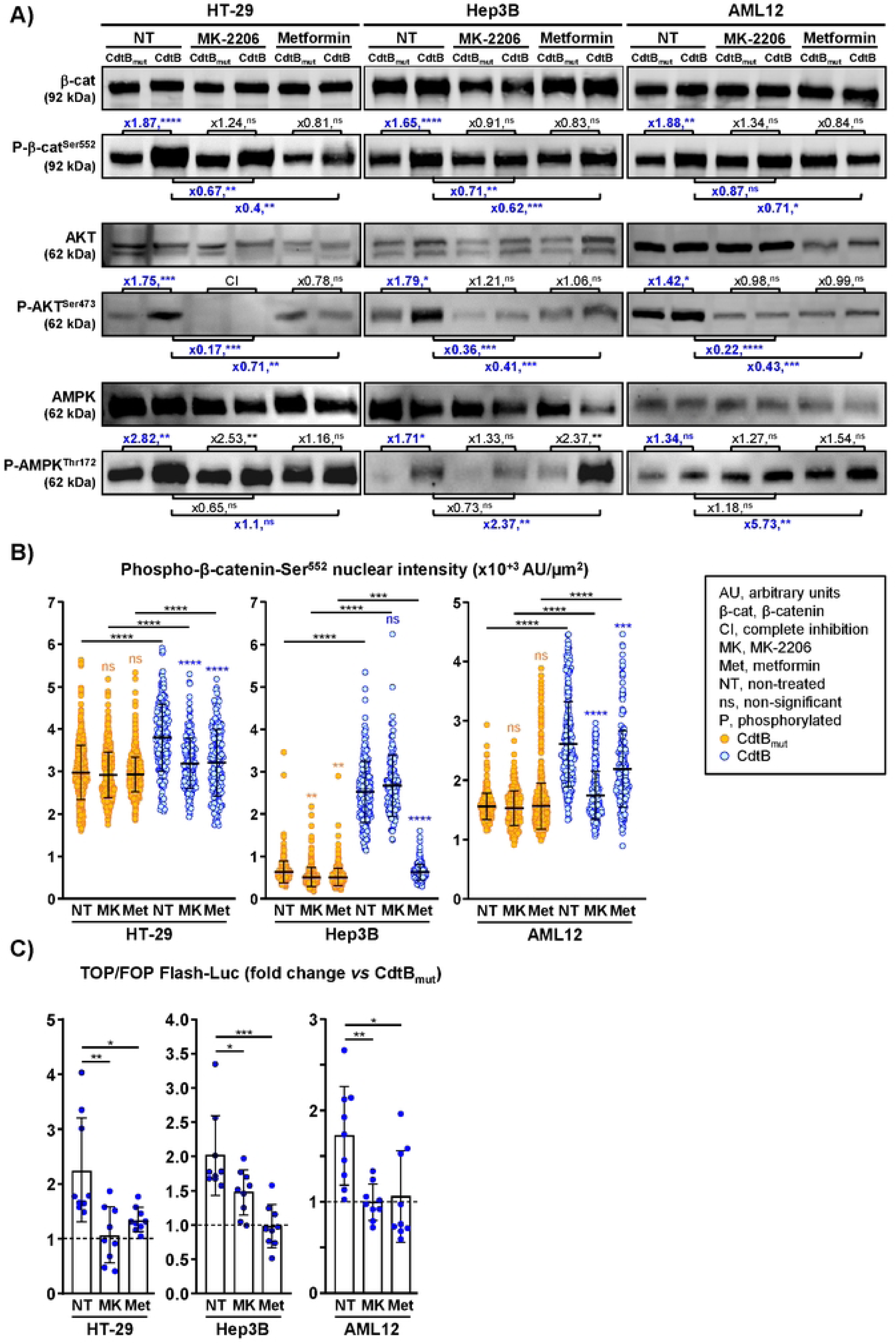
MK-2206 and metformin reduce CdtB-mediated β-catenin Ser552 phosphorylation and nuclear translocation. Transgenic cells were cultured in the presence of doxycycline to induce the expression of CdtB or CdtB_mut_, with MK-2206 (2 µM) or metformin (HT-29 and Hep3B: 5 mM; AML12: 1 mM) added for the final 48 hours prior to analysis. The analyses were conducted 96 hours after induction. **A)** Western blot analysis was performed to examine proteins and their phosphorylated forms. Quantification was conducted relative to total protein levels, using stain-free blots for normalization. Data are presented as the mean ± SD (n=4-6). *p<0.05, **p<0.01, ***p<0.001, and ****p<0.0001 relative to CdtB_mut_ (t-test). Bold blue text represents the fold changes of interest. Metformin failed to further increase CdtB-induced phospho-AMPK-Thr172 in HT-29 cells, indicating CdtB exposure already caused maximal activation. **B)** Cells were probed with primary antibodies generated against β-catenin (red), phosphorylated β-catenin at Ser552 (green) and counterstained with DAPI to highlight the nuclei (blue) prior quantification. Corresponding images are presented in Fig. S7. The nuclear intensity of β-catenin phosphorylated at Ser552 was quantified. Data are presented as the mean ± SD (n=3, >300 nuclei). ***p<0.001 and ****p<0.0001 relative to untreated CdtB_mut_ cells (orange asterisks), or CdtB cells (blue asterisks), and also CdtB relative to CdtB_mut_ for each treatment (black asterisks) (Kruskal-Wallis and Dunn’s post-hoc test). **C)** TOPflash/FOPflash-luciferase reporter activity. Results are presented as means ± SD of 3 independent experiments, each performed in triplicate (n=9). *p<0.05 and ****p<0.0001 (ANOVA and Tamhane’s T2 post-hoc test). *Abbreviations*: AU, arbitrary units; β-cat, β-catenin; CdtB, cytolethal distending toxin active subunit B of *H. hepaticus* strain 3B1; CdtB_mut_, mutated CdtB lacking catalytic activity; ns, non-significant; P-AKT^Ser473^, AKT phosphorylated at Ser^473^; P-AMPK^Thr172^, AMPK phosphorylated at Thr172; P-β-cat^Ser552^, β-catenin phosphorylated at Ser552.

Overall, MK-2206 completely inhibited AKT phosphorylation in HT-29 cells and markedly reduced its levels in Hep3B and AML12 cells. The compound significantly attenuated the CdtB-induced increase in phospho-AKT-Ser473 levels in HT29, Hep3B, and AML12 cells (by 0.17-, 0.36-, and 0.22-fold, respectively). As a result, this led to a reduction in the CdtB-induced increase in phospho-β-catenin-Ser552 levels in HT-29 and Hep3B cells (by 0.67- and 0.71-fold, respectively), while a trend toward reduction was observed in phospho-β-catenin-Ser552 levels in AML12 cells (Fig. 4A).

As expected, metformin treatment strongly increased CdtB-induced increase in phospho-AMPK-Thr172 in Hep3B (2.37-fold) and AML12 (5.73-fold) cells. This effect was not observed in HT-29 cells, suggesting that maximal activation had already been achieved during CdtB exposure. Metformin treatment attenuated the CdtB-induced increase in phospho-AKT-Ser473 levels in HT-29, Hep3B, and AML12 cells (by 0.71-, 0.41-, and 0.43-fold, respectively), confirming the inhibition of CdtB-mediated AKT activation. Consequently, this resulted in a reduction of the CdtB-induced increase in phospho-β-catenin-Ser552 levels in HT-29, Hep3B, and AML12 cells (by 0.4-, 0.62-, and 0.71-fold, respectively).

Treatment with MK-2206 and metformin did not alter overall nuclear levels of phospho-β-catenin-Ser552 in cells exposed to CdtB_mut_, except in Hep3B cells (Fig. 4B and S7). Both treatments, however, reduced the CdtB-induced increase in nuclear phospho-β-catenin-Ser552 levels—except for MK-2206 in Hep3B cells. Despite this inconsistency in the Hep3B cell line, both MK-2206 and metformin attenuated the CdtB-induced enhancement of β-catenin/TCF/LEF-mediated transcriptional activity in HT-29, Hep3B, and AML12 cells (Fig. 4C).

Overall, these findings demonstrate that MK-2206 and metformin suppress the CdtB-induced activation of β-catenin—a process driven by AKT signaling—thereby reducing the CdtB-enhanced β-catenin/TCF/LEF-mediated transcriptional activity.

### MK-2206 and metformin attenuate CdtB-induced EMT features

EMT is characterized by the loss of epithelial features, including the disruption of adherens junctions. β-catenin acts as a crucial molecular switch in this process; its detachment from cell-to-cell junctions and subsequent nuclear translocation drive the expression of genes that promote cell adhesion remodeling, migration, and invasion. MK-2206 and metformin treatment did not affect adherens junction integrity in these cells exposed to CdtB_mut_. Consistent with previous findings, the quantification of cells with junctions could not be performed in Hep3B cells, as cell-cell junctions in these cells exhibit weak cohesion [15]. Exposure to CdtB induced the disassembly and removal of β-catenin from adherens junctions (Fig. 1, S1, 5A,B, S7) in HT-29 and AML12 cells. MK-2206 and metformin treatment reduced the disassembly and removal of adherens junctions in HT-29 and AML12 cells exposed to CdtB (Fig. 5A,B).

**Figure 5.**
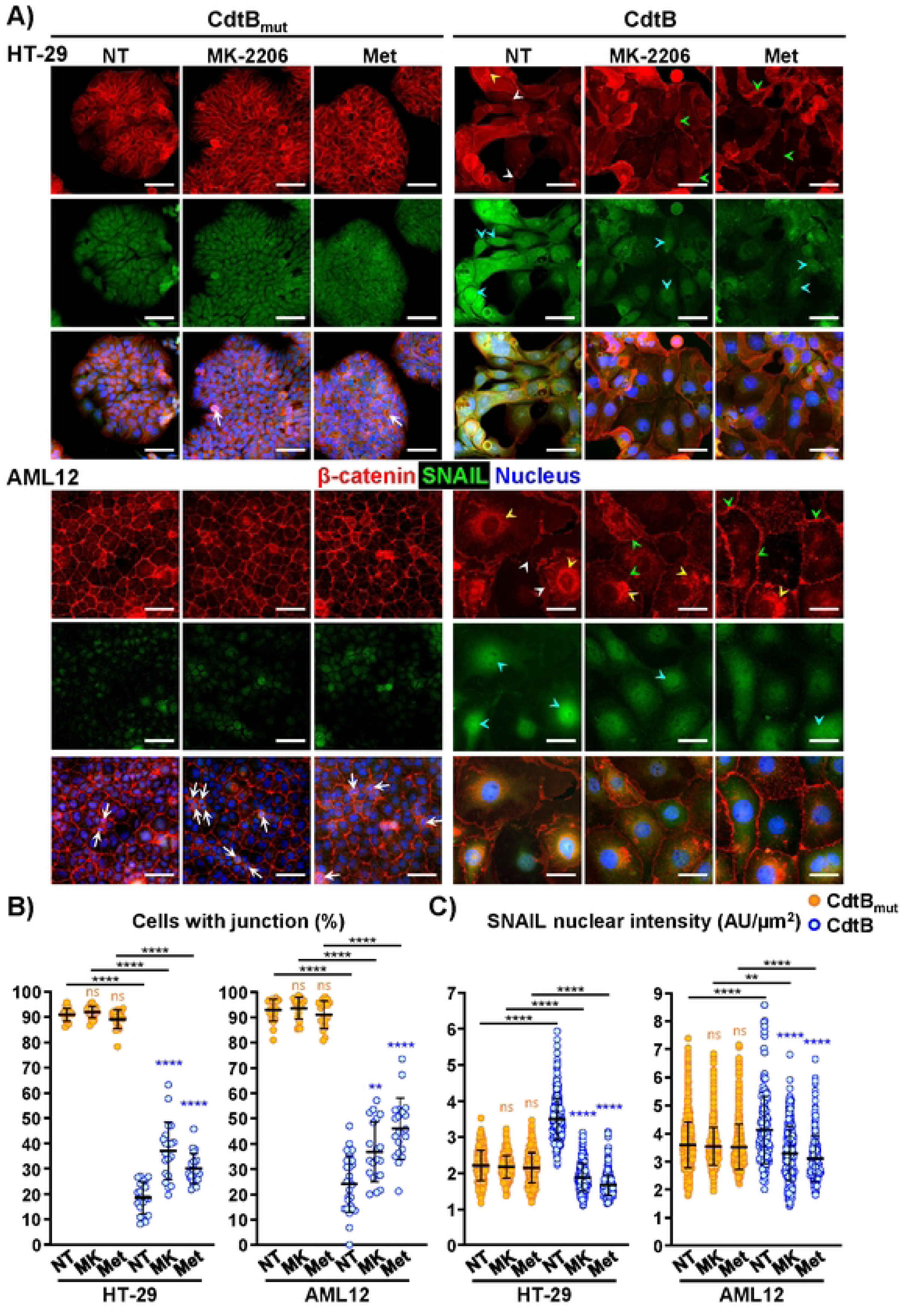
MK-2206 and metformin reduce CdtB-induced cell-cell junction disruption and nuclear accumulation of SNAIL. Transgenic cells were cultured in the presence of doxycycline to induce the expression of CdtB or CdtB_mut_, with MK-2206 (2 µM) or metformin (HT-29: 5 mM; AML12: 1 mM) added for the final 48 hours prior to analysis. At 96 hours post-induction, cells were probed with primary antibodies generated against cell-cell junctions and SNAIL followed by fluorescently labeled secondary antibodies (β-catenin: red; SNAIL: green) and counterstained with DAPI to highlight the nuclei (blue). **A)** Images of cells are presented. Scale bar, 50 μm. White arrowheads indicate the loss of junctional β-catenin and residual membrane spans containing β-catenin. Yellow arrowheads indicate cytoplasmic accumulation and perinuclear relocalization of β-catenin. Blue arrowheads indicate nuclear accumulation of SNAIL. Green arrowheads indicate a lesser loss of junctions in the presence of MK-2206 and metformin (more dense and marked junctions). White long arrows (in merge) indicate bright, condensed chromatin in cells undergoing mitosis. **B)** Cells displaying adherens cell-cell junctions were manually counted across three independent experiments; each using three coverslips divided into three sections. Data represent the mean ± SD (n=27). **p<0.01 and ****p<0.0001 for CdtB_mut_ relative to untreated CdtB_mut_ cells (orange asterisks), CdtB relative to untreated CdtB cells (blue asterisks), and also CdtB relative to CdtB_mut_ for each treatment (black asterisks) (ANOVA and Tamhane’s T2 post-hoc test). The quantification was not performed in Hep3B cells, as cell-cell junctions in these cells exhibit weak cohesion [15]. **C)** Quantification of nuclear SNAIL accumulation. Data represent the mean ± SD (n=3, >300 nuclei). **p<0.01 and ****p<0.0001 for CdtB_mut_ relative to untreated CdtB_mut_ cells (orange asterisks), CdtB relative to untreated CdtB cells (blue asterisks), and also CdtB relative to CdtB_mut_ for each treatment (black asterisks) (Kruskal-Wallis and Dunn’s post-hoc test). Quantification and images of Hep3B nuclear SNAIL accumulation are presented in Fig. S9. *Abbreviations:* AU, arbitrary units; CdtB, cytolethal distending toxin active subunit B of *H. hepaticus* strain 3B1; CdtB_mut_, mutated CdtB lacking catalytic activity; DAPI, 4ʹ,6-diamidino-2-phenylindole; MK, MK-2206; Met, metformin; NT, non-treated; ns, not significant.

The SNAIL transcription factor plays a crucial role in EMT by promoting cell motility [34,35]. In line with our previous findings [15], exposure to CdtB induced a greater nuclear accumulation of SNAIL in HT-29 and Hep3B cells compared to those exposed to CdtB_mut_; however, this increase was less significant in AML12 cells (Fig. 5A,C, S8A, S9). This finding was further confirmed by confocal analysis (Fig. S8A,B). The presence of MK-2206 and metformin did not affect SNAIL nuclear accumulation in cells exposed to inactive CdtB_mut_ (Fig. 5A,C and S9). Treatment with MK-2206 and metformin reduced CdtB-induced nuclear SNAIL accumulation by approximately 2-fold in HT-29 cells, 2.1-fold in Hep3B cells, and 1.3-fold in AML12 cells (Fig. 5A,C and S9). This is consistent with AKT inhibition, which has been shown to decrease nuclear SNAIL levels [35].

During EMT, the enhanced production of MMP drives ECM remodeling, enabling cells to bypass tissue barriers and migrate. We have previously reported increased expression levels of MMPs mRNAs and activities in response to CdtB [15]. To definitively assess the functional consequences of this proteolytic signaling, we performed matrix degradation assays (Fig. 6A). Baseline FITC-gelatin degradation was minimal in AML12 cells exposed to inactive CdtB_mut_, whereas HT-29 and Caco-2 cells exhibited substantially higher basal degradation levels, consistent with the invasive potential of these cancer-derived cell lines. Upon exposure to active CdtB, all cell lines demonstrated significantly elevated proteolytic activity compared to those exposed to CdtB_mut_, showing increases of 13.6-fold in HT-29, 5.8-fold in Hep3B, and 10.6-fold in AML12. MK-2206 treatment doubled matrix degradation in Hep3B cells exposed to CdtB_mut_; this increase likely masked any potential inhibition of CdtB-mediated matrix degradation. MK-2206 treatment did not significantly affect matrix degradation in HT-29 and AML12 cells exposed to CdtB_mut_. However, it reduced CdtB-induced matrix degradation by half in both cell lines.

**Figure 6.**
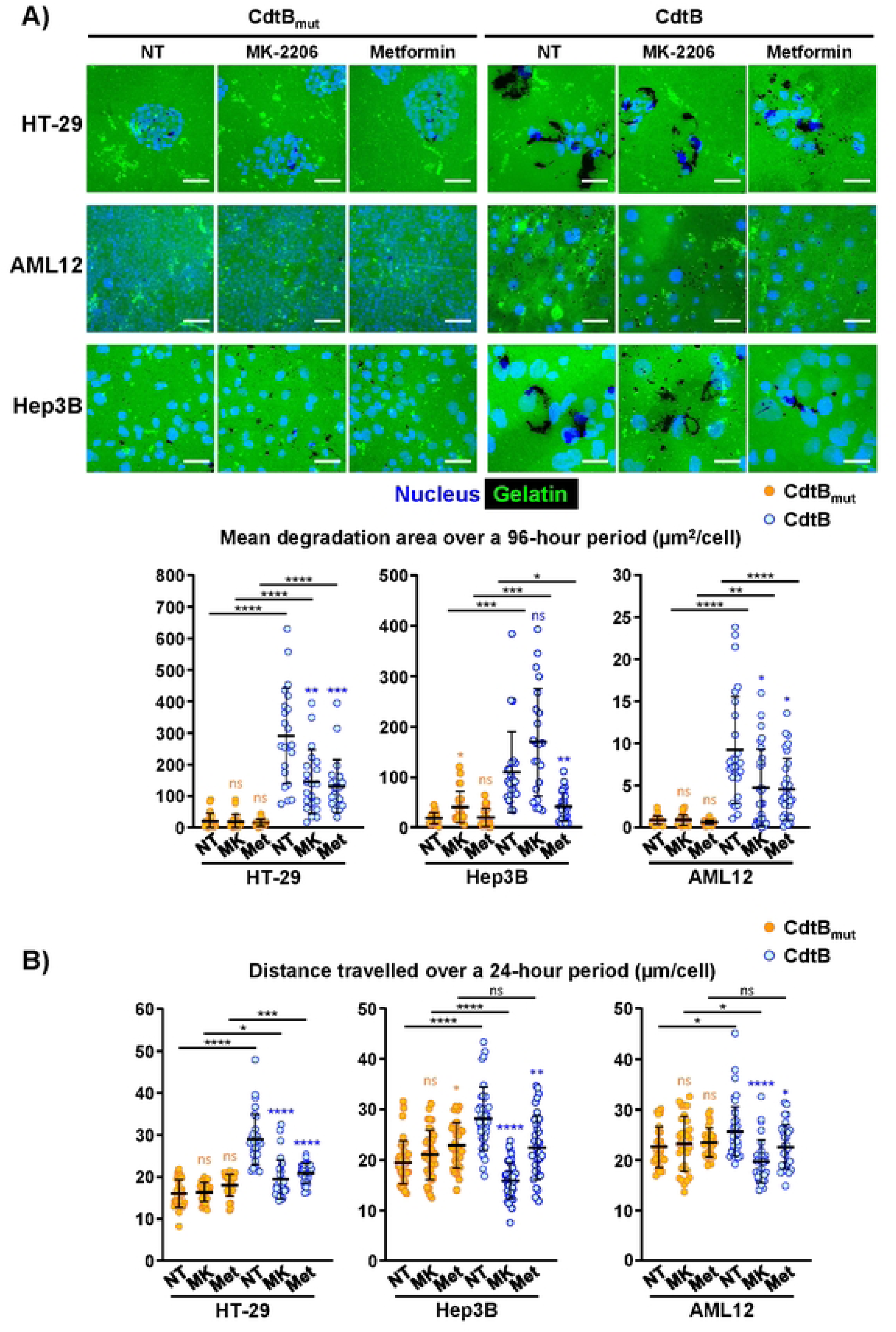
MK-2206 and metformin reduce CdtB-induced matrix degradation and motility. **A)** Transgenic cells were seeded onto gelatin-Oregon Green™ 488 conjugate and cultured in the presence of doxycycline to induce the expression of CdtB or CdtB_mut_, with MK-2206 (2 µM) or metformin (HT-29 and Hep3B: 5 mM; AML12: 1 mM) added for the final 48 hours prior to analysis. At 96 hours post-induction, gelatin degradation was quantified (n=3, >21 fields). **B)** Transgenic cells were seeded without coverslips, treated as described in panel A, and monitored at 30-minute intervals using time-lapse microscopy from 72 to 96 hours post-induction. The graph represents the quantification of the distance travelled by the cells ± SD (n=3, >30 cells, Mann-Whitney U test). Images were captured from the same fields of view for the last 24 hours using an Echo Revolution automated hybrid upright and inverted fluorescence microscope. Cellular motility was measured and expressed as an average distance travelled by the cells per 24 hours (n=30 cells). *p<0.05, **p<0.01, ***p<0.001 and ****p<0.0001 for CdtB_mut_ relative to untreated CdtB_mut_ cells (orange asterisks), CdtB relative to untreated CdtB cells (blue asterisks), and also CdtB relative to CdtB_mut_ for each treatment (black asterisks) (ANOVA and Tamhane’s T2 post-hoc test). *Abbreviations:* CdtB, cytolethal distending toxin active subunit B of *H. hepaticus* strain 3B1; CdtB_mut_, mutated CdtB lacking catalytic activity; DAPI, 4ʹ,6-diamidino-2-phenylindole; FITC, fluorescein isothiocyanate, MK, MK-2206; Met, metformin; NT, non-treated; ns, not significant.

Metformin treatment did not affect matrix degradation in HT-29, Hep3B, or AML12 cells exposed to CdtB_mut_. However, it reduced the CdtB-induced increase in matrix degradation by 2-fold in both HT-29 and AML12 cells, and by 2.6-fold in Hep3B cells. While matrix degradation was monitored over 96 hours of CdtB exposure, MK-2206 and metformin were only administered during the final 48 hours. Consequently, the observed inhibitory effects on CdtB-induced matrix degradation are likely underestimated, given that CdtB-mediated degradation had already progressed for 48 hours prior to their administration.

CdtB-exposed cells undergo EMT, resulting in enhanced migratory capacity [15]. To evaluate the functional role of AKT in this CdtB-induced motility, cells were treated with active or inactive CdtB, with MK-2206 or metformin added during the final 48 hours of incubation. Individual cell trajectories was tracked *via* time-lapse microscopy during the final 24 hours of incubation, providing a precise quantification of the total distance traveled (Fig. 6B). This analysis confirmed that cells exposed to CdtB exhibited increased motility, compared to those exposed to CdtB_mut_, with increases of 1.8-fold in HT-29, 1.5-fold in Hep3B and 1.2-fold in AML12. Metformin treatment slightly enhanced motility in Hep3B cells exposed to CdtB_mut_, which may have attenuated its inhibitory effect on CdtB-induced motility, although the inhibition remained significant. Consistent patterns of motility inhibition by MK-2206 and metformin across all cell lines suggest a common underlying mechanism (Fig. 6B).

Despite minor cell-line variations—specifically, MK-2206’s lack of effect on nuclear phosphorylated β-catenin (Ser552) accumulation or matrix degradation in Hep3B cells—both MK-2206 and metformin consistently suppressed CdtB-induced β-catenin/TCF/LEF transcriptional activity. Furthermore, both inhibitors effectively attenuated downstream EMT features, including cell–cell junction disruption, SNAIL nuclear accumulation, matrix degradation, and cell motility

In conclusion (Fig.7), CDT/CdtB intoxication induce the phosphorylation of β-catenin at Ser552, along with its loss from adherens junctions. This triggers the subsequent cytoplasmic accumulation and nuclear translocation of β-catenin, ultimately driving TCF/LEF-mediated transcription—a pivotal downstream event of Wnt/β-catenin signaling. Critically, pharmacological blockade of AKT-mediated phosphorylation of β-catenin at Ser552 prevented these effects and reduced CdtB-induced SNAIL nuclear accumulation, matrix degradation, and cellular motility.

**Figure 7.**
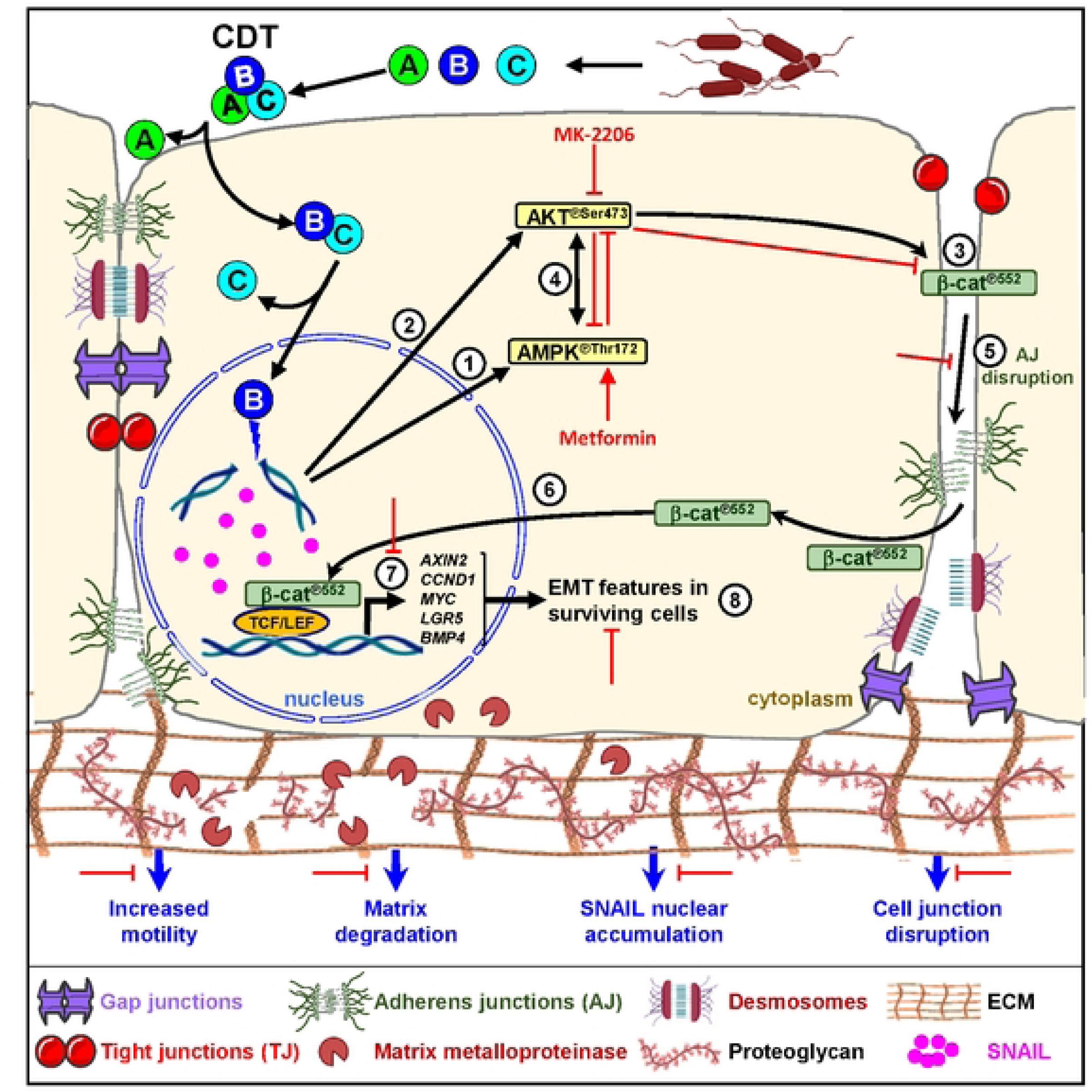
CdtB controls epithelial-to-mesenchymal transition through β-catenin dependent signaling. The CDT heterotrimeric complex (CdtA, CdtB, and CdtC) is secreted by Gram-negative bacteria. CdtA and CdtC allow the internalization of the CdtB-CdtC heterodimer into the host cell, while CdtA remains anchored to the cell surface. Within the cell, the CdtC subunit undergoes degradation in the endoplasmic reticulum, and CdtB is transported to the nucleus, where it induces DNA damage. CdtB exposure promotes the phosphorylation of both ① AMP-activated protein kinase (AMPK) and ② AKT, resulting in ③ β-catenin phosphorylation at Ser552. Under the effect of CdtB, the phosphorylation of AKT activates that of AMPK and *vice versa* ④. Phosphorylation of β-catenin at Ser552 promotes ⑤ its dissociation from cell-cell junctions, leading to the disruption of adherens junctions. This, in turn, releases free phosphorylated β-catenin into the cytosol, where it is ⑥ subsequently translocated to the nucleus to enhance TCF/LEF-mediated transcriptional programs⑦. These effects, observed in surviving cells, mediate features of ⑧ the epithelial-mesenchymal transition. Specifically, inhibiting the phosphorylation of β-catenin at residue 552 by selective (MK-2206) and non-selective (metformin) inhibitor of AKT reduces CdtB-induced disruption of cell-cell junctions and nuclear accumulation of phosphorylated β-catenin, thereby reducing β-catenin-TCF/LEF transcription. Furthermore, this inhibition also diminishes CdtB-induced nuclear accumulation of SNAIL, matrix degradation, and cellular motility. *Abbreviations:* CDT, cytolethal distending toxin; Ⓐ, CdtA; Ⓑ, CdtB; Ⓒ, CdtC; ECM, extracellular matrix; EMT, epithelial-mesenchymal transition.

## Discussion

Bacterial genotoxins, such as CDT and colibactin, have emerged as significant contributors to cancer development through their ability to induce DNA damage in host cells [36]. Chronic exposure to these toxins, produced by certain pathogenic bacteria, causes DNA double-strand breaks. This is not merely a cytotoxic event; it also interferes with cellular processes, promoting genomic instability through an impaired DNA damage response and the acquisition of malignant phenotypic properties. This includes profound actin cytoskeleton remodeling [23] and EMT [15], which are fundamental drivers in carcinogenesis.

This study provides compelling evidence that these genotoxins manipulate the Wnt/β-catenin signaling pathway which, when dysregulated, plays a critical role in the development and progression of various cancers and other diseases [37]. β-catenin has dual functions: it serves as a structural component of adherens junctions and as a co-transcription factor that drives TCF/LEF-mediated transcription of Wnt/β-catenin target genes. Our findings demonstrate that CDT, specifically through its active subunit CdtB, promotes the phosphorylation of β-catenin at Ser552, leading to the loss of β-catenin from adherens junctions in both non-transformed cells and cancer-derived cells. This phosphorylation event is crucial as it stabilizes β-catenin and facilitates its nuclear translocation. In the nucleus, β-catenin interacts with TCF/LEF transcription factors, driving TCF/LEF transcription, a key downstream event in Wnt/β-catenin signaling. This cascade of events suggests a mechanism through which CDT/CdtB can promote oncogenic transformation and exacerbate cancer in patients already diagnosed with the disease, by dysregulating Wnt/β-catenin signaling and enhancing TCF/LEF transcription.

Colibactin, a complex secondary genotoxic metabolite produced by *E. coli,* also causes the removal of β-catenin from adherens junctions and its activation, leading to TCF/LEF-mediated transcription. Colibactin indirectly damages DNA, notably through the formation of crosslinks [38], whereas CdtB is a DNAse that directly cleaves DNA [39]. Despite their distinct structures and mechanisms of action, both toxins induce DNA double-strand breaks [38,39] and exhibit some similar effects, such as of cell-cell junction disruption and enhancement of TCF/LEF-mediated transcription, as shown in this study. This suggests that the observed effects on β-catenin signaling are primarily triggered by the induced DNA damage. This was confirmed using DNA-damaging agents (etoposide and streptozocin), which also activate β-catenin-TCF/LEF-mediated transcription. Taken together, these findings suggest that the genotoxic stress induced by CDT and colibactin is likely the primary factor initiating the loss of cell-cell junctions and the activation of Wnt/β-catenin signaling.

CDT- and colibactin-induced modulation of β-catenin occurs in a cellular context marked by persistent DNA damage. Both toxins activate DNA damage response pathways, enforcing long-term G2/M arrest, endoreplication-induced polyploidy, and senescence—features reflected in the enlarged, flattened morphology of intoxicated cells with persistent DNA damage foci [1–3,24,36,38–40]. In this setting, β-catenin activation links genotoxin-induced genomic instability to transcriptional programs that promote both damaged cell survival and mesenchymal trait acquisition. This connection between persistent genotoxic stress, β-catenin activation, and EMT aligns with evidence that Wnt/β-catenin signaling is engaged downstream of DNA damage to drive adaptation, stemness, and therapy resistance in cancer cells [41]. Senescent cells, metabolically active and secreting a senescence-associated secretory phenotype (SASP), synergize with β-catenin activation and SNAIL-dependent EMT to remodel the microenvironment, thereby fostering the emergence and dissemination of more aggressive, neighboring non-senescent cell clones [42,43]. Supporting this, CDT- and colibactin-induced junctional loss of β-catenin may weaken contact inhibition, enhances paracrine signaling, and modulates crosstalk with EMT- and SASP-central pathways [5,24,44,45]. Critically, the loss of junctional β-catenin coupled to its transcriptional activation, indicate that these toxins reprogram epithelial signaling networks rather than merely disrupting junctions. Given the persistence of senescent cells and their SASP secretion, CDT-driven senescence—combined with β-catenin-dependent EMT and matrix degradation—may establish a pro-tumorigenic microenvironment that fosters clonal selection of genotoxin-damaged clones. This framework supports the dual role of bacterial genotoxins as both initiators and promoters of carcinogenesis, simultaneously inducing DNA damage, disrupting cell cycle control, and hijacking β-catenin signaling to enable the survival and dissemination of genetically unstable cells.

The differential response to CdtB in cancer (HT-29, Hep3B) and non-cancer (AML12) cells suggests that the toxin exploits specific vulnerabilities in cancer cells, resulting in more pronounced effects compared to non-transformed cells, where β-catenin regulation is tightly controlled. One hypothesis could be that cancer cells, due to their altered genetic and epigenetic landscapes, have dysregulated pathways that govern cell-cell junctions, β-catenin activation, SNAIL nuclear translocation, extracellular matrix degradation, and motility. These pathways might be more sensitive to CdtB-induced stress, resulting in enhanced disruption and activation. Conversely, non-cancer cells may have intact regulatory mechanisms that make them more resilient to the toxin, thereby maintaining their cellular integrity and function more efficiently.

β-catenin undergoes numerous post-translational modifications, including phosphorylation at 58 distinct sites reported to date [46]. Phosphorylation at the N-terminal leads to its degradation by the proteasome, whereas phosphorylation at Ser191/605 by JNK2 [18], Ser552 by AKT [19] and Ser675 by PKA[20], stabilizes β-catenin. AKT-mediated phosphorylation of β-catenin at Ser552 is a key step in regulating its cellular localization and function [19]. The use of MK-2206, a selective AKT inhibitor, provided insights into the signaling pathways involved in CDT/CdtB-induced effects within the population of cells surviving exposure to both the toxin and the inhibitor. Overall, MK-2206 strongly diminished AKT activation and protected both non-transformed and cancer-derived cells from several effects induced by CdtB exposure, including the disassembly of cell-cell junctions, the accumulation of phosphorylated β-catenin at Ser552 (notably in the nucleus), and the increased TCF/LEF-mediated transcription. However, the inhibition of β-catenin phosphorylation at Ser552 by MK-2206 remained incomplete, particularly in HT-29 cells where AKT inhibition was fully achieved. This suggests that other kinase(s) may be directly or indirectly involved in this phosphorylation process. For example, PKA, known to phosphorylate β-catenin at Ser675, can mediate β-catenin phosphorylation at Ser552 [47,48]. A role for PAK1/2 (P21 RAC1 Activated Kinase 1/2) [49], PGK1 (Phosphoglycerate Kinase 1) [50], TBK1 (TANK Binding Kinase 1) [51], CAMK2A (Calcium/Calmodulin Dependent Protein Kinase II Alpha) and MSK1 (RPS6KA5, Ribosomal Protein S6 Kinase A5) [52] was also recently reported. AMPK was reported to directly phosphorylate β-catenin at Ser552, thereby enhancing TCF/LEF-mediated transcription [53]. However, it is unlikely that the increase in activated AMPK observed upon CdtB exposure is the primary mechanism for β-catenin activation. This conclusion is supported by the observation that metformin efficiently activated AMPK in cells exposed to inactive and active CdtB, but also led to a reduction in CdtB-induced β-catenin phosphorylation and activation.

Activated AMPK can negatively regulate the PI3K/AKT pathway by attenuating PI3K activity, which, particularly in cancer cells, reduces AKT phosphorylation at Ser473 [54]. Therefore, in cells exposed to CdtB, metformin treatment significantly enhances AMPK activation, leading to reduced AKT phosphorylation at Ser473 and ultimately decreasing β-catenin phosphorylation at Ser552. These findings underscore the central role of AKT signaling in mediating cellular responses to CDT/CdtB.

β-catenin, AKT, and EMT are deeply intertwined molecular players that collectively drive various aspects of cancer initiation, progression, metastasis, and therapy resistance [55]. Their dysregulation forms a complex network that promotes the aggressive behavior of many tumor types. Chronic/persistent activation of the Wnt/β-catenin and PI3K/AKT signaling pathway often cross-talk to cooperatively promote EMT in various cellular processes, including cancer progression [55]. Direct inhibition of AKT activation (MK-2206) protected against several CdtB-induced effects associated with EMT, such as the disruption of cell-cell junctions, increased nuclear accumulation of SNAIL, enhanced matrix degradation, and higher cellular motility. Metformin treatment conferred similar protection against CdtB exposure, thereby confirming its ability to reverse EMT [56–58].

Metformin acts on pathways targeted by CDT and colibactin, such as β-catenin and AMPK signaling pathway, which are often associated with carcinogenesis. Metformin, a biguanide derivative, is widely prescribed as first-line pharmacotherapy for type 2 diabetes mellitus. It inhibits mitochondrial complex I (NADH dehydrogenase), increasing the phosphorylation of AMPK at Thr172 [59]; it also inhibits the Wnt/β-catenin signaling pathway [60], which is involved in the pathogenesis of CRC. This antidiabetic drug is associated with a reduced incidence of cancer and cancer-related mortality among diabetic patients, notably digestive cancers [61–63]. It shows significant promise in preventing genomic instability, DNA repair enhancement, and oxidative stress reduction [64]. Extensive clinical trials are currently underway worldwide, investigating the repurposing of this antidiabetic drug for the prevention and treatment of various cancers, including gastrointestinal cancers. Our findings support metformin as a pertinent inhibitor of the Wnt/β-catenin signaling pathway and highlight its efficiency as a very promising neoadjuvant to be combined with conventional anticancer therapies. This is particularly relevant for colorectal cancer patients, who are frequently infected with genotoxin-producing bacteria.

In conclusion, this study underscores the critical role of bacterial genotoxins —specifically CDT— in dysregulating β-catenin signaling, a process that drives EMT and ECM degradation, ultimately remodeling the microenvironment. These alterations may foster a supportive niche that enhances cancer initiation and progression. Notably, our findings reveal that AKT inhibition could mitigate the pro-carcinogenic effects of these genotoxins, aligning with the observed cancer-preventive potential of metformin—particularly in gastrointestinal cancers, where CDT and colibactin are frequently implicated. Collectively, these results position these genotoxins at the intersection of DNA damage responses, cell cycle regulation, β-catenin signaling, and EMT, offering a mechanistic framework for how chronic exposure to genotoxin-producing bacteria may induce both genomic instability and early invasive behavior in epithelial tissues.

## Materials & Methods

### Other materials and methods

The antibodies, compounds, and their concentrations used are listed in Table S1. Supplementary Materials and Methods comprise the ethical statement; protocols for mouse infection, euthanasia, and embedding of liver specimens; cell lines and culture conditions description, techniques for immunodetection, confocal microscopy, image analysis, and protein quantification; Western blot analysis, including image analysis and protein quantification; and RT-qPCR experimental protocols and gene expression quantification.

Loss of cell-cell junctions and other EMT phenotypes emerge 72 hours post-genotoxin exposure [15], whereas significant β-catenin TCF/LEF-mediated transcription is only observed at 96 hours (Fig. 3C). Consequently, all subsequent analyses were conducted at this 96-hour time point.

### Bacteria and infection experiments

*Helicobacter hepaticus* 3B1/Hh-1 and its corresponding isogenic CDT mutant strain with a disrupted *cdtABC* coding region [65] and lacking CDT activity were cultivated as previously reported [66]. The suspensions were prepared in Brucella broth with an OD_600_ adjusted to 1 corresponding to a concentration of 3.3×10^8^ colony forming units (CFU)/ml.

*E. coli* strain DH10B hosting the bacterial artificial chromosome (BAC) vector (*pks-*) or the BAC pks island encoding colibactin (*pks+*) were cultivated as previously reported [21]. The suspensions were prepared in Luria Bertani medium with an OD_600_ adjusted to 1.0 corresponding to a concentration of 5×10^8^ CFU/ml.

For infection experiments, cells were seeded and, the following day, bacteria were added into the medium at a multiplicity of infection (MOI) of 100. The analyses were conducted 96 hours after the onset of infection. For *H. hepaticus*, incubation proceeded for 96 hours without further medium changes, as this bacillus does not survive beyond a few hours in these conditions. In contrast, *E. coli* cocultures were incubated for 4 h, after which the medium was replaced with fresh medium supplemented with gentamicin (10 μg/ml) to inhibit further bacterial growth. Incubation then continued for 92 hours. It should be noted that the HT-29 cell line was not used for infection experiments with *H. hepaticus*, as these cells do not show CDT effects during infection [23].

### Transgenic cell lines

AML12, HT-29, Caco-2 and Hep3B-derived transgenic *cdtB* and mutated *cdtBH_155_L-H_265_L* cell lines were established by lentiviral transduction. Briefly, the pTRIPz lentiviral plasmid with two independent promoters was used: the UBC promoter allowed the constitutive expression of the gene for resistance to puromycin, and the tetracycline response element (TRE) promoter was inducible by tetracycline. The complete *cdtB* sequences of *H. hepaticus* (from the start codon until the codon proximal to STOP codon, GenBank accession numbers: AE017125 or AF163667) fused at their 3ʹ end to three repeats of the influenza hemagglutinin epitope (HA) (GenBank accession numbers: KT590046 and KT590047) were cloned downstream of the TRE promoter in this plasmid instead of the TurboRFP gene initially present. Cells having the integrated transgene sequence in a transcriptionally silent form were selected in the presence of puromycin (2 μg/ml). When required, the transgene expression was induced in the cells from the tetracycline-inducible promoter by addition of doxycycline (600 ng/ml) to the culture medium. The analyses were conducted 96 hours after induction.

### Ethic statement

Animal material provided from previous studies was approved by the Ethics Committee for Animal Care and Experimentation CEEA 50 in Bordeaux (Comité d’éthique en matière d’expérimentation Animale agréé par le ministre chargé de la Recherche, “saisine” 13126B 1, Bordeaux, France), according to treaty no. 123 of the European Convention for the Protection of Vertebrate Animals. All animal experiments were performed in security level 2 animal facilities at the University of Bordeaux by trained authorized personnel only.

### Analysis of β-catenin-mediated TCF/LEF transcription using TOPFlash/FOPFlash Dual Luciferase Reporter Assay

Cells were cultured in 24-well plates for 24 hours. Cells were transfected using Lipofectamine 2000 (#11668019, Thermo Fisher Scientific, France) with 10 ng/well of control pRL-SV40 Renilla luciferase plasmid (an internal control plasmid for normalization, #E2231 Promega) and 200 ng/well of either TOPFlash or FOPFlash reporter plasmids which contain TCF binding sites or mutated TCF sites upstream of a Firefly luciferase reporter gene [29] in Opti-MEM medium without antibiotics and serum. 6 hours later, the cells underwent various treatments for a period of 24-96 hours. Then, cells were lysed with passive lysis buffer (Promega, #E194). Firefly and Renilla luciferase luminescence were measured sequentially from the same sample using the Dual-Luciferase® Reporter Assay System (#E1980, Promega). Firefly luciferase activity was first measured by adding firefly Luciferase Assay Reagent (LARII, #E1980) in cell lysate; 10 sec later, Renilla luciferase activity was measured by adding Stop & Glo reagent (Promega, #E1980). Luminescence was measured sequentially with Lumat LB 9507 luminometer (Berthold®, Gallardon, France). TOPFlash activity was calculated as the ratio of pTOPFlash (firefly/renilla) to the mean of pFOPFlash (firefly/renilla).

### Degradation assay

Transgenic *cdtB* and *cdtB_mut_* cells were seeded on glass coverslips in 24-well plates with FITC-gelatin (1 mg/ml) for 24 h. Then, cells were cultured in the presence of doxycycline to induce the expression of the active CdtB or CdtB_mut_, with MK-2206 or metformin added for the final 48 h prior analysis. At 96 hours post-induction, cells were washed sequentially in PBS for 5 min each, followed by a final staining with 4’,6-diamidino-2-phenylindole (DAPI). A 24 x 60 mm microscope cover glass was placed on top and sealed with nail polish. Degraded fluorescent-free surface was selected and measured using the ‘Threshold’ function of Fiji (v1.53j) software [67].

### Motility assay

Cells were treated as described in ‘Degradation assay’, without coverslips. For the final 24 hours of treatment, time-lapse images were captured hourly from the same fields of view using an Echo Revolution automated hybrid upright and inverted fluorescence microscope (Echo, San Diego, CA) at 10x magnification. Individual cell movement was then measured using the Manual Tracking plugin in Fiji (v1.53j) software [67].

### Statistical analysis

Statistical analyses were performed using GraphPad Prism version 8.0.2 (GraphPad software, San Diego, CA). Outliers were not removed. Data are presented as the mean ± standard deviation. All data sets were considered independent. The choice of test was based on the normality of the data, assessed within the “Normality and Lognormality test” function of GraphPad Prism. The F-test was employed to compare variances between the two datasets. Student’s t-test (with or without Welch’s correction), or Mann-Whitney U test, was selected as appropriate, based on the normality and homogeneity of variances. For comparisons involving more than two independent data sets, the Brown-Forsythe test (for data not normally distributed and sample size less than 15-20) or Bartlett’s test (for other data) was used to check for equal variances. Kruskal-Wallis test with Dunn’s multiple comparisons test, one-way ANOVA with Tukey’s multiple comparisons test, or Welch’s ANOVA with Games-Howell’s multiple comparisons test (n>50/group) or Tamhane T2 test (n<50/group) were selected as appropriate, based on the normality and homogeneity of variances.

## Acknowledgments

This work was supported, in part, by Ligue Contre le Cancer, Gironde (France) and the Ministère de la Recherche et de la Technologie, INSERM and Université de Bordeaux (UMR1312). We are indebted to James G. Fox (Massachusetts Institute of Technology, Cambridge, MA, USA) for supplying the *H. hepaticus* strains. We wish to thank Eric Oswald (Institut de recherche en santé digestive INRA, Toulouse, France) for supplying the *E. coli* strains. We would like to thank Camille Tocqueville and Christophe Grosset (BoRdeaux Institute of onCology-BRIC UMR1312, INSERM, Team ‘MIRCADE, Methods and Innovations for the Research in Pediatric Cancers’) for supplying the plasmids used for TOPFlash/FOPFlash Dual Luciferase Reporter Assay. Confocal microscopy was performed at the Bordeaux Imaging Center, a core facility of CNRS, INSERM, and the University of Bordeaux, and a member of the national infrastructure France BioImaging (ANR-24-INBS-0005 FBI BIOGEN), supported by the French National Research Agency. We acknowledge the assistance of Mónica Fernández-Monreal.

## Presented in part

European workshop on bacterial protein toxins-ETOX. June 29 - July 3, 2025, Oñati, Guipuzkoa, Spain. Armelle Ménard (invited speaker).

Max von Pettenkofer-Institut, Ludwig-Maximilians-University, Munich, München, Germany, October 16, 2025, (Armelle Ménard, invited speaker).

## Data availability statement

All data are available in manuscript and supplementary file.

All raw data were available on 4TU.ResearchData (dataset ‘CDT hijacks β-catenin signaling to promote EMT’, https://data.4tu.nl/private_datasets/R0TiyfE2nDIt-EZ0JB-VI3-vYwyKNCUxtmVHAyqCtFY

## Generative Artificial Intelligence

The Generative AI tool Mistral developed by Mistral AI (https://mistral.ai/) was utilized for proofreading of the text.

